# Diversity and biogeography of Woesearchaeota: A comprehensive analysis of multi-environment data

**DOI:** 10.1101/2020.08.09.243345

**Authors:** Jing Xiao, Yu Zhang, Wanning Chen, Yanbing Xu, Rui Zhao, Liwen Tao, Yuanqi Wu, Yida Zhang, Xiang Xiao, Ruixin Zhu

## Abstract

Woesearchaeota is a newly proposed archaeal phylum frequently detected in various environments. Due to the limited systematical study, little is known about their distribution, taxonomy, and metabolism. Here, we conducted a comprehensive study for Woesearchaeota with 16S ribosomal RNA (rRNA) gene sequencing data of 27,709 samples and metagenomic whole genome sequencing (WGS) data of 1,266 samples. We find that apart from free-living environments, Woesearchaeota also widely distribute in host-associated environments. And host-associated environmental parameters greatly affect their distribution. 81 Woesearchaeota genomes, including 33 genomes firstly reconstructed in this project, were assigned to 59 Woesearchaeota species, suggesting their high taxonomic diversity. Comparative analysis indicated that Woesearchaeota have an open pan-genome with small core genome. Metabolic reconstruction showed that particular metabolic pathway absence in specific environments, demonstrated the metabolic diversity of Woesearchaeota varies in differences environments. These results have placed host-associated environments into the global biogeography of Woesearchaeota and have demonstrated their genomic diversity for future investigation of adaptive evolution.

## 1. Introduction

In the past few years, an increasing number of archaeal phyla have been proposed, which have greatly deepened our understanding for the ecological and evolutionary roles of Archaea domain^1-5^. Woesearchaeota is an archaeal phylum proposed by Castelle *et al*. based on metagenomic analysis^6^. With the limited genomic data extracted from environmental samples, Woesearcheota is considered as one of the most widely distributed archaea in DPANN superphylum^7, 8^. They have been detected in sediments, groundwater, soil, deep-sea hydrothermal vents, hypersaline lakes, wetland, permafrost, and human lung^6, 9, 10^. However, how the geochemical settings define the distribution pattern of Woesearcheaota and their ecological function, especially on a global scale, remain unclear.

Moreover, members within Woesearchaeota phylum appear to have highly divergent, sometimes deficient, metabolic potentials, and this further hinders the identification and isolation of Woesearchaeota. For example, based on 16S rRNA gene sequences, 26 potential subgroups were detected, although the taxonomic documentation was ambiguous^9^. Reconstruction of metabolic pathways from metagenome-assembled genomes (MAGs) of Woesearchaeota showed the absence of certain core biosynthesis, suggesting a symbiotic or parasitic lifestyle^6, 7, 9^. If this deficiency of independent living is a common phenomenon in the entire phylum, it would be particularly intriguing in the evolutionary point of view. Considering the lack of Woesearcheota isolate, comparative genomics study based on large datasets of WGS should be a promising approach to access the genomic and metabolic feature of Woesearchaeota.

To date, with the accomplishment of several world-class microbiome projects, such as Earth Microbiome Project (EMP) and *Tara* Oceans Project^11-14^, more data from various environments are available for a systematically investigation. To expand knowledge for distribution, taxonomy, and metabolism of Woesearchaeota, we conducted a comprehensive study with combination of two types of data. 16S rRNA gene sequencing data were used to explore their distribution characteristic, meanwhile, metagenomic data were collected to investigate taxonomic and metabolic diversity of Woesearchaeota. This endeavor allowed us to construct a framework for distribution, taxonomy, and metabolism of Woesearchaeota, contributing to guidance to efficient cultivation and profound investigation.

## 2. Results

### 2.1 Biogeography of Woesearchaeota

#### Widely distribution of Woesearchaeota

16S rRNA gene sequences from EMP were collected to explore the distribution characteristics of Woesearchaeota. A total of 27,709 samples were carefully analyzed in this study, and we got 23,428 qualified samples, among which 6,788 samples were identified as containing Woesearchaeota. These Woesearchaeota are widely distributed around the world, in both marine and inland environments (Fig. 1A). Further analysis revealed that Woesearchaeota not only present in free-living environments such as water, soil, and sediments, but also live in host-associated environments such as plant rhizosphere, biofilm, animal surface, and animal secretion (Fig. 1B). Apparently, Woesearchaeota are more often to be discovered in free-living environments rather than in host-associated environments. In free-living environments, Woesearchaeota are most widely distributed in the sediment, while in host-associated environment, Woesearchaeota are most extensively distributed in rhizosphere.

**Fig. 1.**
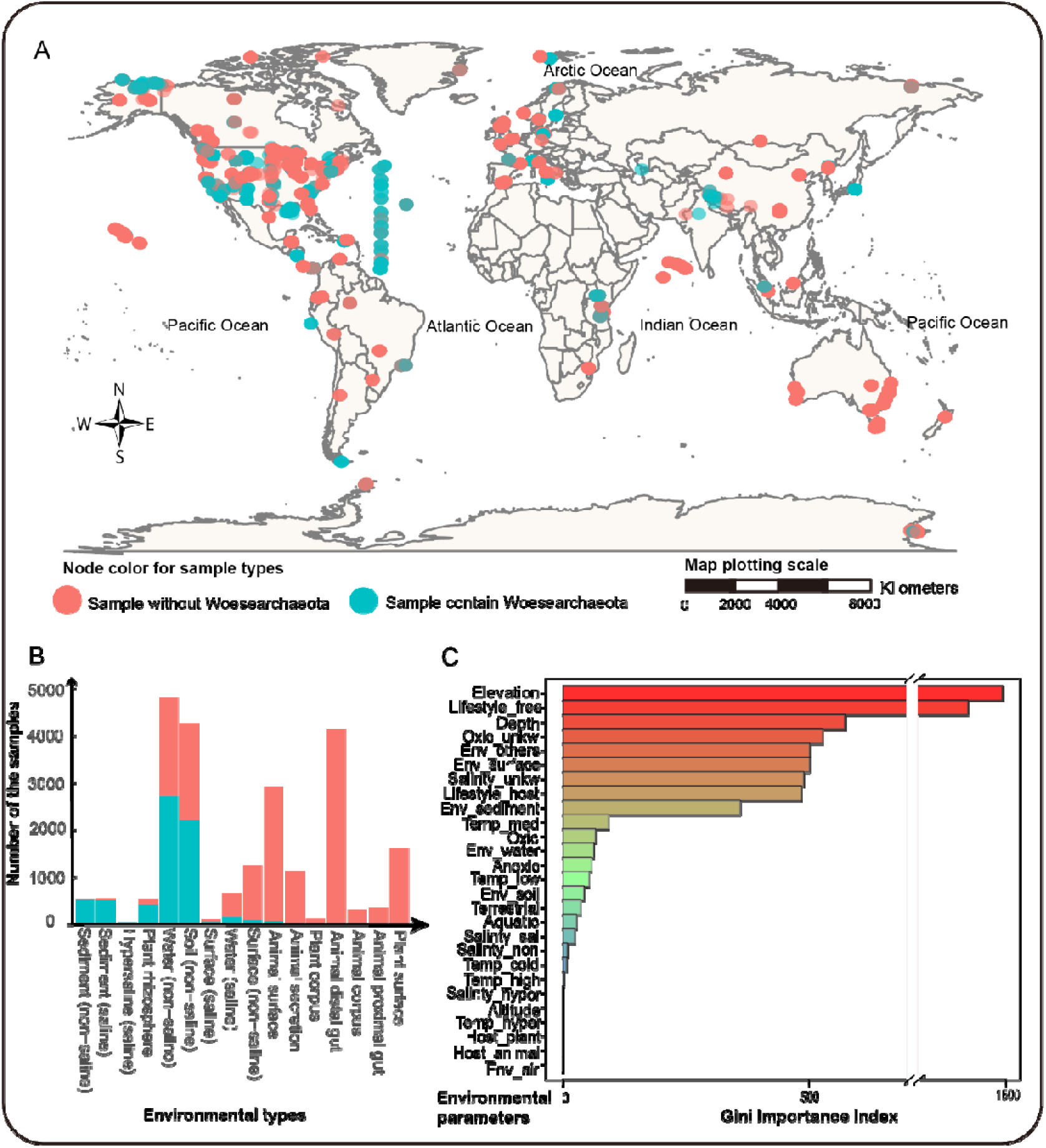
Distribution characteristics of Woesearchaeota. **A**. Global distribution of EMP samples showed the present/absence of Woesearchaeota. **B**. Distribution of Woesearchaeota in different types of environment (*Number of samples from hypersaline environment is relatively small compared to other environments, for all 13 samples, 10 of them contain Woesearchaeota). **C**. Importance of different environmental factors affecting the distribution for Woesearchaeota.

#### Impact of environmental parameters

To assess how environmental parameters effect the distribution of Woesearchaeota, Random Forest classifier model ^15^ was constructed, and 27 environmental parameters were taken as input feature vectors. The mean area under the curve (AUC) values of the model is 0.9467(10-fold cross validation). Feature importance is evaluated by Gini index (Fig. 1C). Among all the environmental factors, elevation and depth of the samples are of great importance, affecting the distribution of Woesearchaeota. Besides, 6 features (Lifestyle_free, Oxic_unkw, Env_others, Salinity_unkw, Lifestyle_host, Temp-med) related to free-living/host-associated lifestyle also matters, which consistent with the finding that Woesearchaeota are mainly distributed in free-living environments. For the remaining environment factors, oxygen conditions, temperature, terrestrial/aquatic, and salinity show a decreasing importance, but they also affect the distribution of Woesearchaeota. While altitude, host type (plant/animal) almost have no effect on their distribution.

### 2.2 Taxonomy of Woesearchaeota

#### Genome collection of Woesearchaeota

To explore the taxonomic characteristics of Woesearchaeota, genomes of Woesearchaeota in public database were also collected. We got 48 high-quality Woesearchaeota genomes (Supplementary Fig.1) after de-duplication and quality control for all 105 candidate genomes of Woesearchaeota. Among all high-quality genomes, nearly 90% of the genomes were from water sample including groundwater, marine water, and hot spring water, while only 4 genomes from sediments and 1 genome from soil.

#### Genome reconstruction and phylogeny of Woesearchaeota

We collected over ∼35 terabyte metagenomic WGS data from dominant habitats of Woesearchaeota, including samples from sea water, rhizosphere and sediment. After trimmed all these data, de novo assembly and binning were then conducted, resulted in the reconstruction of 74 high quality (>70% completeness and <5% contamination) target archaeal genome bins. Phylogeny based on 16 concatenated ribosomal proteins reveals that these archaeal bins belong to different archaeal clades (Fig. 2). And 33 genome bins belong to Woesearchaeota, among which 5 from sediments, 23 from sea water, and 5 from host-associated environment rhizosphere (Supplementary Table 1).

**Table 1.** Genomic information of 17 nearly-complete Woesearchaeotal genomes.

| No | Genome Id | Data Source | Completeness | Contamination | Biosample | Metagenome_source |
| --- | --- | --- | --- | --- | --- | --- |
| 01 | GCA_005222965.1 | NCBI | 99.18 | 1.65 | SAMN07236660 | marine sediment |
| 02 | GCA_002867475.1 | NCBI | 98.9 | 2.2 | SAMN07982670 | enrichment from estuary sediment |
| 03 | GCA_000830295.1 | NCBI | 98.9 | 1.1 | SAMN03202995 | Rifle groundwater |
| 04 | GCA_002688315.1 | NCBI | 98.63 | 0 | SAMN07982670 | enrichment from estuary sediment |
| 05 | GCA_002762785.1 | NCBI | 97.8 | 0 | SAMN06659469 | Groundwater ( 2m ) |
| 06 | ERR594345.116 | In house | 97.53 | 0 | SAMEA2619974 | sea water |
| 07 | ERR599062.136 | In house | 97.53 | 1.1 | SAMEA2619970 | sea water |
| 08 | ERR594345.48 | In house | 96.98 | 2.2 | SAMEA2619974 | sea water |
| 09 | SRR8464959.7 | In house | 96.89 | 0 | SAMN10350739 | Carex aquatilis--Peat Soil |
| 10 | GCA_002762985.1 | NCBI | 96.7 | 1.1 | SAMN06659468 | Groundwater ( 2m ) |
| 11 | EPRCH7.83 | In house | 96.7 | 1.1 | OEX003658 | Black smokers |
| 12 | ERR594345.135 | In house | 96.43 | 1.1 | SAMEA2619974 | sea water |
| 13 | GCA_003695265.1 | NCBI | 96.15 | 1.1 | SAMN10119972 | hot springs metagenome |
| 14 | ERR594345.31 | In house | 96.15 | 2.2 | SAMEA2619974 | sea water |
| 15 | SRR8464960.7 | In house | 95.97 | 0 | SAMN10350571 | Carex aquatilis--Peat Soil |
| 16 | GCA_002498125.1 | NCBI | 95.6 | 1.1 | SAMN06027228 | soil metagenome |
| 17 | EPRCH7.420 | In house | 95.6 | 1.1 | OEX003658 | Black smokers |

**Fig. 2.**
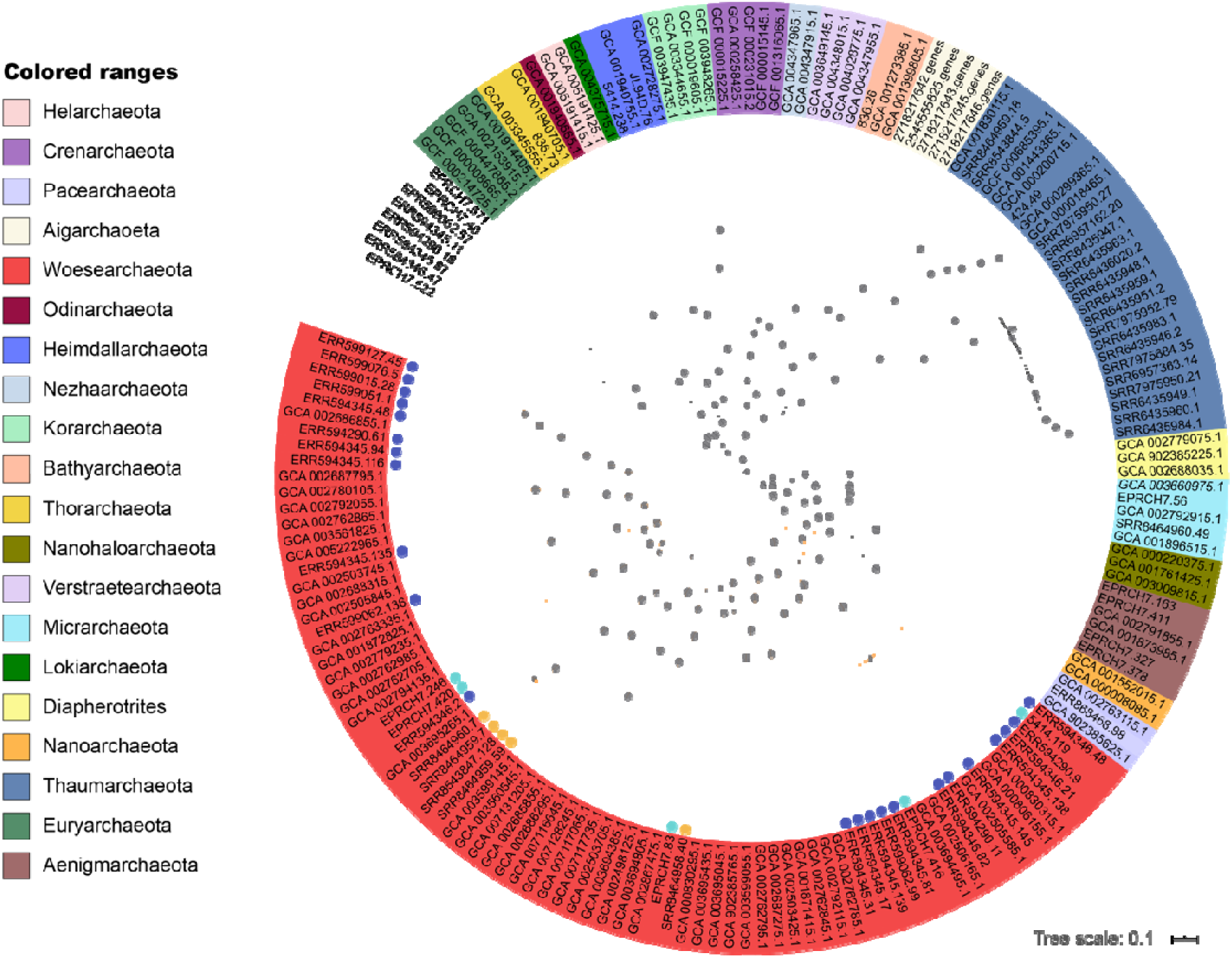
Phylogeny of archaeal phyla. Each leaf represents an archaeal genome, grey dot on each branch represents bootstrap value, ranging from 0.6 to 1. and red range represent Woesearchaeota, leaves with colored dot mean the Woesearchaeotal genomes reconstructed in this study, different color represents different sample source. (dark blue dot: *Tara* Oceans Project; light blue dot: in-house data; orange dot: NCBI-SRA).

#### Taxonomic groups of Woesearchaeota

For further investigation of Woesearchaeotal taxonomic characteristics, 48 high-quality genomes of Woesearchaeota from public database were also used. Adding up 33 high-quality Woesearchaeota genome bins reconstructed in this study, we finally gathered 81 high-quality Woesearchaeota genomes for further study(Supplementary Fig.1). CheckM tool was used to evaluate genome quality, and among all these high-quality genomes bins, more than half of the genome bins have completeness higher than 90%. Moreover, all these Woesearchaeota genome bins are of small size (averagely 1.04 Mb), encoding 1,174 genes on average.

To accurately identify the taxonomic groups of Woesearchaeota genomes, we used whole genome sequences. Pairwise orthoANI (ANI, Average nucleotide identity)^16, 17^ calculation among all 81 genomes were conducted. Based on previous studies, orthoANI value takes a similar range of cut cut-off as ANI for species demarcation, which is approximately 95–96%^18^. The taxonomic identification results showed that all 81 genomes belong to 59 Woesearchaeal species (Fig. 3), and orthoANI values among most genome bins are ∼63%, showing that the Woesearchaeota are of high taxonomic diversity at the species level. Meanwhile, 19 new species of Woesearchaeota have been discovered in our study, including 2 species first discovered from host-associated environments.

**Fig. 3.**
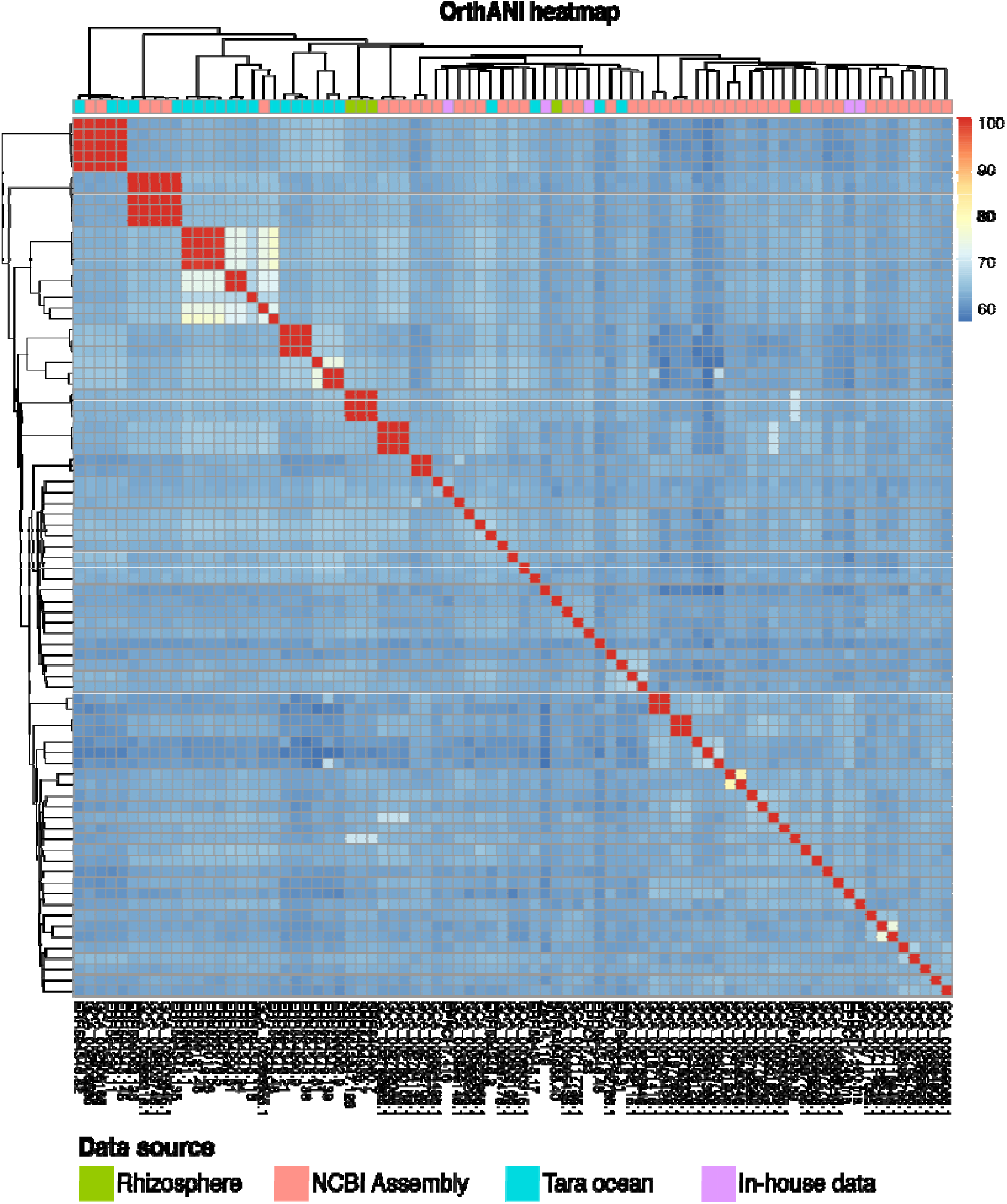
Taxonomic diversity of Woesearchaeota. Each grid represents the OrthoANI value between two corresponding genomes.

### 2.3 Open pan-genome with limited core genome genes

Among all 81 high-quality Woesearchaeota genomes, 17 genome bins (Table 1; Supplementary Fig.1) are nearly-complete (completeness > 95%) with relatively low contamination (contamination < 2.2%). Thus, a comparative genomics anlysis for Woesearchaeota was conducted by using these genomes. A total of 20,731 predicted protein-coding-genes were obtained, which were clustered into 15,109 orthologous clusters. The power-law regression analyses indicated an open pan-genome for Woesearchaeota (Supplementary Fig.2). Besides, the contributions of core, accessory, and unique genes in Woesearchaeal pan-genome (Fig. 4 A) showed that they only contain a small core genome. On average, only 3.2% of genes in each Woesearchaeal genome are core genes, and the rest are accessory genes and unique genes, accounting for 47.3% and 49.5% genes in each genome, respectively. Moreover, proportion of accessory genes and unique genes varies in different Woesearchaeal genome, and the percentage of unique genes is higher than accessory genes in most genomes.

**Fig. 4.**
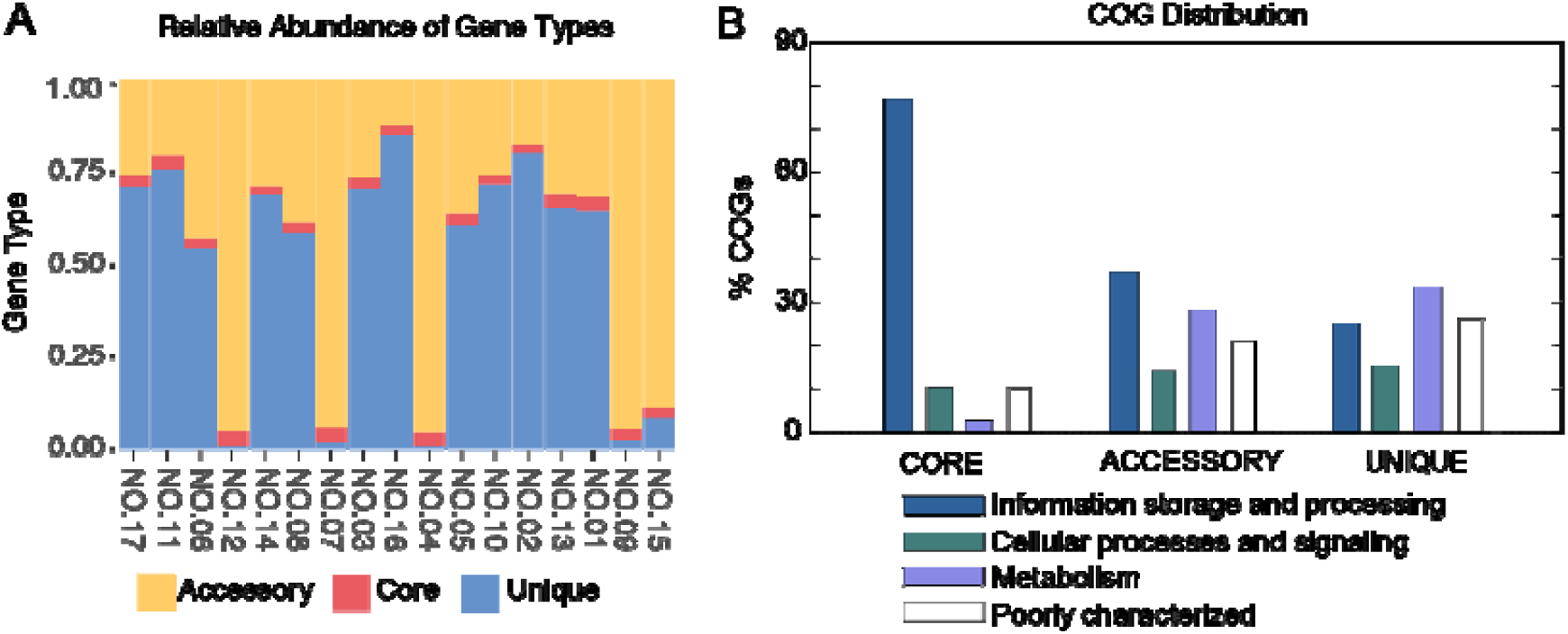
Constituent and function of Woesearchaeotal Pan-genome. **A**. Relative abundance of core/accessory/unique genes in Woesearchaeotal genomes. **B**. Distribution of genes function.

Further investigation revealed function profiles of Woesearchaeal pan-genome (Fig. 4 B). For core genes, most are assigned to “information storage and processing”, followed by “cellular processes and signaling”, which only make up a much smaller proportion. Compared to core genes, differences of function profiles are relatively small in accessory genes and unique genes. And the majority of accessory genes and unique genes are assigned to “information storage and processing” and “metabolism”, respectively. Meanwhile, poorly characterized genes presented in all three groups, and functions of these genes are still not clear, over 25% of the unique genes are so-called “poorly characterized” genes.

### 2.4 Metabolic capabilities of Woesearchaeota

To explore their metabolic capability, we used 17 nearly-complete genomes (Table 1) for further analysis. Metabolic reconstruction showed that most of the core biosynthetic pathways are incomplete in Woesearchaeota. For example, none of the Woesearchaeal genomes encodes the complete tricarboxylic acid cycle (TCA cycle). And most of the genes encoding respiratory-associated enzymes are absent in all these genomes. Besides, in a large proportion of these Woesearchaeal genomes, glycolytic pathway is incomplete because of the absence of few genes (Fig. 5).

**Fig. 5.**
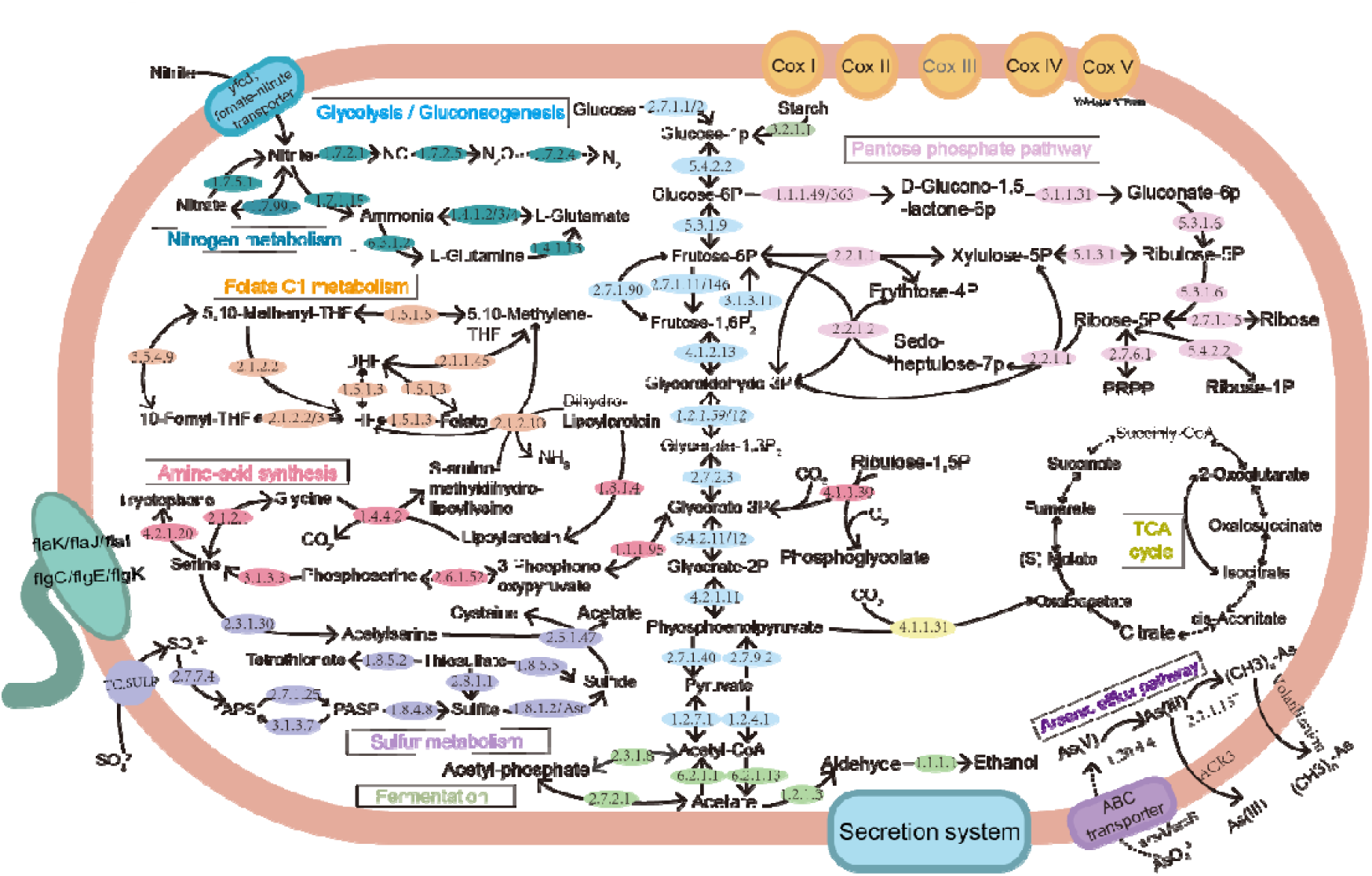
Metabolic pathways of Woesearchaeota. Pathways were constructed based on KEGG database, the soiled arrows mean related genes were presented in some genomes while the dotted arrows mean absence of corresponding genes in all genomes.

#### Carbon metabolism

A complete glycolytic pathway in Woesearcheaota was first discovered in this study. In previous studies, the gene *pfk* encoding phosphofructokinase was absent in all the Woesearchaeotal genomes. However, this gene was found in several genomes with the accumulation of nearly-complete Woesearchaeotal genomes. It can be inferred that most Woesearchaeota can convert glucose. Notably, we discovered gene *porA/B* in some Woesearchaeotal genomes, which has not been reported before. The *porA/B* gene encodes enzyme converting pyruvate to acetyl-CoA, meanwhile, some other Woesearchaeota accomplish the conversion by encoding pyruvate dehydrogenase.

#### Nitrogen metabolism

Dissimilatory nitrate reduction pathway was first found in Woesearchaeota. And *narG* and *nirD* genes are discovered in Woesearchaeotal genome, encoding nitrate reductase and nitrite reductase respectively. These enzymes enable the transformation of Nitrite to Ammonia. Moreover, genes encoding enzymes catalyzing denitrification were also detected, including *narG, nirK*, and *norC* genes, while *nosZ* gene was not discovered in our study.

#### Sulfur metabolism

Only Assimilatory sulfate reduction pathways presented in Woesearchaeotal genomes. In these genomes, sulfate is reduced to APS (Adenylyl sulfate) firstly, then reduced to PAPS (3’-Phosphoadenylyl sulfate). Afterwards, *cysH* gene encodes the enzyme catalyzing PAPS to sulfite. Finally, either gene (*ars*/*cysJ* gene) encode enzyme to reduce sulfite to sulfide. *Ars* and *cysJ* genes present in different Woesearchaeotal genomes, encoding anaerobic sulfite reductase and sulfite reductase respectively. Additionally, we also found other sulfur metabolism related genes, including *doxD, TST, phsA, cysK*, and *cysE*.

#### Arsenic metabolism

Interestingly, several genes involved in arsenic metabolism were discovered in some Woesearchaeotal genomes for the first time, such as *arsC2* and *ACR3* genes. In these genome, *arsC2* gene encodes arsenate reductase, which reduces arsenate to arsenite, and then arsenite is pumped out of the cells by using the transporter encoded by *ACR3*. Moreover, *AS3MT* and *arsC* genes also presented in some Woesearchaeotal genomes, indicating arsenic metabolism capability.

### 2.5 Metabolic capability of Woesearchaeota varies in differences environments

Metabolic reconstruction for Woesearchaeota from different environments shows that metabolic capability of Woesearchaeota differs in various environments (Fig. 6). As for nitrogen metabolism, most Woesearchaeotal genomes contain predicted genes for dissimilatory nitrate reduction and denitrification, however, no genes related to nitrogen metabolism presents in the genomes of Woesearchaeota from soil. Meanwhile, metabolic reconstruction shows all Woesearchaeota genomes contain sulfur metabolism related pathways, comparative analysis for their metabolic capability reveals that none of Woesearchaeota from black smokers have genes encoding enzymes that catalyze assimilatory sulfate reduction.

**Fig. 6.**
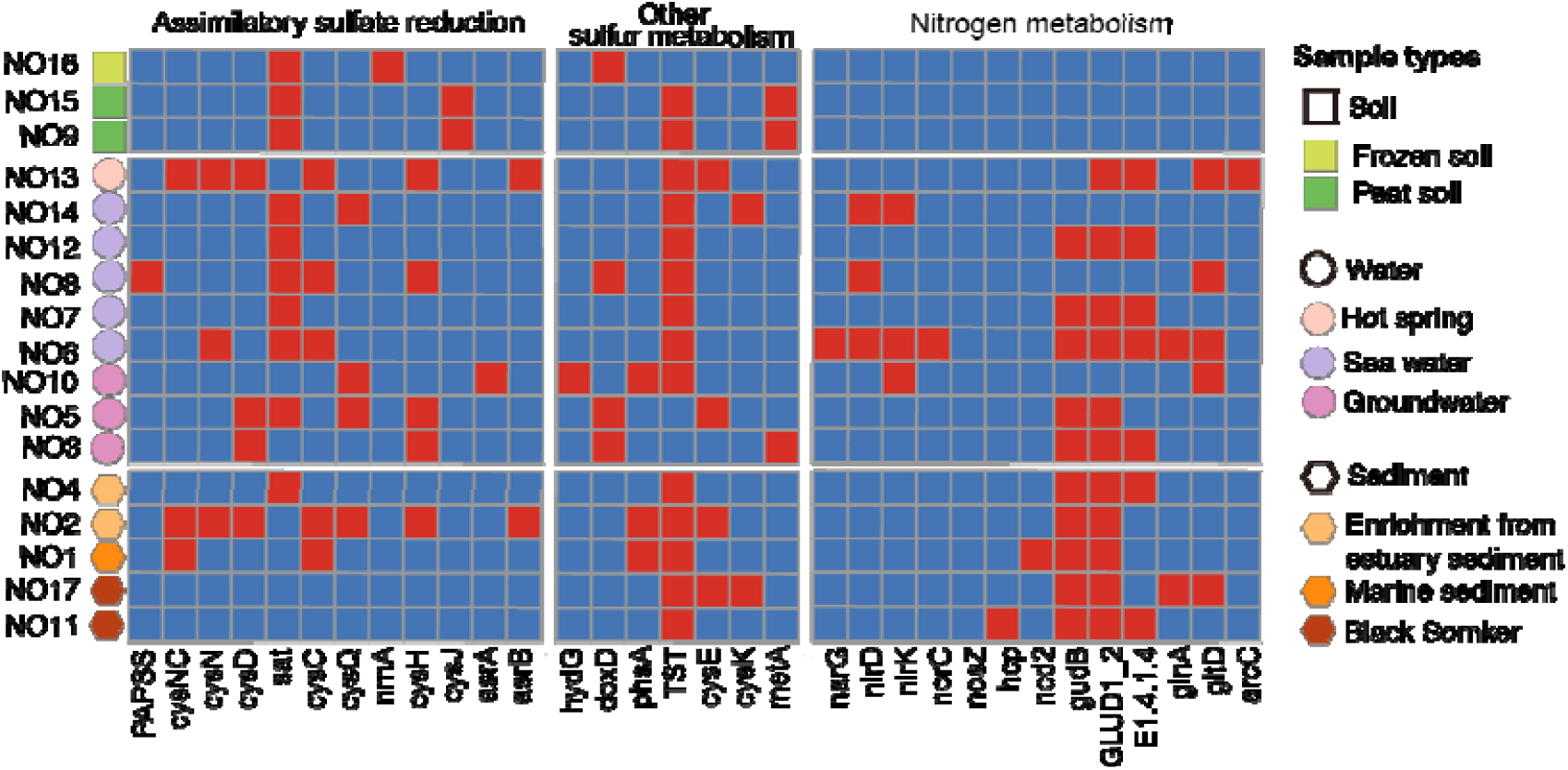
Metabolic capabilities of Woesearchaeota from different environments. Red grid means the gene was detected in corresponding genome, while blue grid indicates the absence of the gene. Numbers on the left represent genome ID in Table 1.

## 3. Discussion

In this study, a comprehensive analysis greatly expanded the knowledge of Woesearchaeotal diversity in aspects of their distribution, taxonomy and metabolism. Distribution analysis based on 16S rRNA gene sequencing data suggested that Woesearchaeota also have a wide distribution in host-associated environments, especially in plant rhizosphere, which greatly expanded our understanding for distribution diversity of Woesearchaeota. The investigation of environmental parameters revealed that apart from elevation and depth of these samples, host-associated environmental parameters also play an important part in the distribution of Woesearchaeota. Besides, oxygen condition is of greatest importance among the remaining environmental parameters, which is consistent with previous analysis^9^. The results suggest that the location rather than the geochemical conditions plays a major role on driving the distribution pattern of Woesearchaeota. Meanwhile, it is important to note that only elevation and depth are consecutive data, while most parameters are discrete data due to the limitation of current data. With accumulation of more environmental parameters, future work may develop a full picture of the distribution characteristic of Woesearchaeota.

Besides, exploration of the taxonomic characteristic revealed that Woesearchaeota are of high taxonomic diversity, with 81 genomes assigned to 59 species. Taxonomic study for Woesearchaeota expanded the phylogenetic diversity of Woesearchaeota by adding up 19 new species of Woesearchaeota, which account for 32% of all Woesearchaeotal species, indicating analyses for metagenomic data from various environments can greatly deepen our understanding for Woesearchaeota, and the discovery of new Woesearchaeotal species will promote further taxonomic study. Moreover, pan-genome studies for Woesearchaeota show that the core genome of Woesearchaeota is small. Considering comparative analysis was on the phylum level, the great distance among all genomes may account for the relatively small core genome. Highly diversity and small core genome of Woesearchaeotal implies that Woesearchaeota have strong ability in speciation, further research focusing on this ability may reveal evolution mechanism of Woesearchaeota, which is also known as fast-evolving archaeal taxa.

Meanwhile, with the extended dataset, Woesearchaeota is confirmed as with small genome sizes with limited metabolic capabilities, suggesting a host-dependent lifestyle. Metabolic reconstruction combined with pan-genome analyses provided a framework to explore the metabolic diversity of Woesearchaeota. Woesearchaeota have a large open pan-genome, of which accessory genes and unique genes make up a large proportion. And over 30% of these genes are assigned to “Metabolism”, suggesting unique metabolic way in different Woesearchaeotal genomes. These genes play an important role in the diversity and adaptability of Woesearchaeota. Further investigation for Woesearchaeota from various environments shows that Metabolic capability of Woesearchaeota differs in different environments. Although some archaea were reported to have a significant impact on nitrogen cycles in soils^19^, the only environment where Woesearchaeota have deficiency in nitrogen metabolism was soil. Meanwhile, Woesearchaeota from black smokers are unable to conduct assimilatory sulfate reduction. It is known that black smokers are rich in sulfur-bearing minerals, they emit particles such as H_2_S and FeS, which provide microorganism with energy by oxidation^20^. Since sulfur plays an important role in this environment, it is vital to investigate whether the lack of specific genes is influenced by the environment. Lateral gene transfer (LGT) is an important driving force in the evolution of microorganisms^21-24^, thus, it could conceivably be hypothesized that Woesearchaeota gained these genes by LGT from the environments to help them adapt to the environments. And future researches should be undertaken for hypothesis testing.

## 4. Conclusions

In summary, we performed a comprehensive study to investigate the distribution, taxonomy, and metabolism of Woesearchaeota. The distribution pattern suggested that Woesearchaeota are widely distributed in various biotopes, including host-associated environments. Then, 33 high-quality Woesearchaeotal genomes were reconstructed from metagenomic data, greatly expanded taxonomic group of Woesearchaeota. And these genomes are collected for further taxonomic study and revealed that Woesearchaeota are of high taxonomic diversity. Meanwhile, Comparative genomic analysis showed that Woesearchaeota have a large open pan-genome with small core genome. Metabolic reconstruction for Woesearchaeota implied their metabolic potential for Nitrogen, Sulfur, and Arsenic. Moreover, metabolic capacity of Woesearchaeota varies in different environments, suggested that they are of high metabolic diversity. This study greatly expanded our knowledge for distribution, taxonomy, and metabolism of Woesearchaeota. And it demonstrated great diversity of this archaeal phylum in different aspects. However, due to the limitation of current data, our knowledge about the driving force of metabolic diversity in Woesearchaeota is limited. Thus, future researches are encouraged to explore this problem.

## 5. Data and methods

### 5.1 Data collection

#### 16S rRNA gene sequencing data

8,023,841 represented Operational taxonomic unit (OTU) sequences from 27,709 samples were collected from EMP^11, 12^. Meanwhile, OTU composition and environmental parameters for all samples were gathered.

#### Woesearchaeotal metagenome-assembled genomes

All metagenome-assembled genomes (MAGs) of Woesearchaeota from public database and previous studies were collected. These genomes were retrieved (by 8th October, 2019) from NCBI Assembly database (https://www.ncbi.nlm.nih.gov/assembly) matching the query string “(“Candidatus Woesearchaeota"[Organism] OR woesearchaeota[All Fields]) AND latest[filter]”. Furthermore, Woesearchaeotal genomes from prior studies were also gathered^6, 9^. Then a custom perl script was used for de-duplication, and CheckM (version 1.0.13) was used to estimate the quality of these genomes. Only high-quality (completeness>70% and contamination<5%) genomes were used for further investigation.

#### Metagenomic WGS data

A total of ∼35 terabyte metagenomic data were collected for metagenomic study, including samples from rhizosphere, sediment, and water. Based on our distribution exploration, among all host-associated environments, Woesearchaeota are most widely distributed in rhizosphere. Thus, metadata for metagenomes of 102 rhizosphere samples collected from NCBI-SRA (https://trace.ncbi.nlm.nih.gov/Traces/sra/). Meanwhile, only few genomes have been reconstructed from sediment samples in public database, thus we used in-house marine sediments samples, including samples of black smoker and marine sediments (see **Data availability** for detail). Moreover, *Tara* Oceans Project provides a systematic sampling data for marine microbe^13, 25^, and 1,158 water samples were collected from this project.

### 5.2 Distribution characteristic exploration

Taxonomic classification was conducted using BLAST+, all OTU sequences were BLASTed against SILVA SSU128^26, 27^. For each OTU sequence, we filtered results with percent identity less than 0.80, and the OTU sequence will be annotated as Woesearchaeota while over 51% of the filtered results belonging to Woesearchaeota^28^. Quality control for all samples (kept samples marked as qc_filtered ==“TRUE” in EMP && counts>10,000 && no missing parameters) with OTU composition analysis identified samples containing Woesearchaeota (relative abundance of Woesearchaeotal OTU > 0.01%).

#### Environmental parameter analysis

We used Random Forest ^15^ to investigate the impact of environmental parameters on the distribution of Woesearchaeota. First, a label is assigned to each sample based on existence/absence of Woesearchaeota. Second, by utilizing the sample environment factors, such as altitude, depth, oxic/anoxic, salinity, etc., a feature vector containing all environmental parameters is generated for each sample. Third, a Random forest classifier is trained and 10-fold cross validation was implied for evaluation (R package ‘randomForest’). Last, Gini importance index are calculated to estimate feature importance^29^.

### 5.3 Genome reconstruction from metagenomic data

For all those metagenomic data, raw sequencing reads are trimmed by using Trimmomatic (version 0.38; SLIDINGWINDOW:10:20 MINLEN:50)^30^. Then trimmed short reads were assembled to long contigs using MEGAHIT (version 1.1.3) with default parameters^31^. Qualified reads were then mapped to contigs using BamM (version 1.7.3; http://ecogenomics.github.io/BamM/) to calculate the coverage information. Afterwards, genome binning was performed on contigs by MetaBAT2 (version 2.12.1) with default parameters^32^, and the minimum size of a contig for binning is 2,500. CheckM (version 1.0.13) was used to estimate the completeness and contamination of all bins^33^.

### 5.4 Taxonomy analysis

For phylogenetic analysis, representative archaeal reference genomes were downloaded from NCBI Assembly database. Then, CheckM was used to estimate the quality of these genomes, and high-quality reference genomes were collected. Meanwhile, target high-quality genome bins (Marker lineage annotated as “k Archaea (UID2)” in CheckM) reconstrued from metagenomic data were also used for phylogenetic analysis. Gene prediction was performed using Prodigal (version 2.6.3) with “-p meta”^34^. HMM models were used to identify 16 ribosomal proteins (L2, L3, L4, L5, L6, L14, L15, L16, L18, L22, L24, S3, S8, S10, S17, S19) from all archaeal genomes using HMMER (version 3.1b2) with “hmmsearch -E 1E-5”^6, 35, 36^. Genomes with less than 8 ribosomal proteins were not included in the analyses. Then, individual proteins were aligned with MUSCLE(version 3.8.3.1)^37^, trimmed using trimAL (version 2.0) with “-automated1”^38^. Maximum-Likelihood phylogeny of 16 concatenated proteins using both fasttree(version 2.1.0; -lg -gamma) and IQ-TREE (version 1.6.12; -st AA -m MFP -bb 1000 -nt 16)^39, 40^.

Woesearchaeotal genomes reconstructed from metagenomic data combined with reference genomes belong to Woesearchaeota were collected for taxonomic identification(Supplementary Fig.3). OrthoANI value were calculated pairwise by using OrthoANI tool (https://www.ezbiocloud.net/tools/orthoani)^18^.

### 5.5 Pan-genome profiling

Taken the reduced genomes of Woesearchaeota in consideration, quality estimation for all Woesearchaeotal genomes were conducted using CheckM with a refined marker set of Archaea. And thoese nearly complete genomes (completeness>95%, contamination <5%) were used for comparative genomics analysis. Protein sequences for each genome were predicted using Prodigal^34^. USEARCH was used for orthologous clustering with 50% sequence identity taken as cut-off value. And the power-law regression model and exponential curve fit model were used to calculated the pan-genome size and core genome size, respectively. Then, we analyzed the distribution of core gene, accessory gene and unique gene in each Woesearchaeotal genome. In addition, function annotation for each orthologous protein cluster is based on protein BLAST against reference COG (Clusters of Orthologous Groups of proteins) and KEGG databases^41, 42^. Protein clustering, pan-genome profile analysis, and function and pathway analysis are conducted using BPGA-pipeline^43^ (https://iicb.res.in/bpga/downloads.html).

### 5.6 Metabolic prediction

Nearly complete genomes of Woesearchaeota were collected to perform metabolic reconstruction. Prodigal was used to predict open reading frames (ORFs) from these genome bins. The ORFs were annotated by using eggnog-mapper(v2)^44, 45^, and resulting data contained Gene Ontology (GO) terms, KEGG Orthology (KO) and archaeal clusters of orthologous genes (arCOGs). KEGG metabolic pathways was reconstructed for each genome by using KO with KEGG mapper tool^42^. To infer metabolic capacities of Woesearchaeota from different environments, environmental factors are combined for a comparative analysis.

## Data availability

Woesearchaeotal high-quality genomes reconstructed in this study have been deposited at NODE (https://www.biosino.org/node/) under accessions OEP000995. Besides, other high-quality genomes reconstructed from *Tara* Oceans Project and rhizosphere samples have been deposited at NODE under accessions OEP000994 and accessions OEP000996, respectively.

Moreover, in-house metagenomic data used in this study have been deposited at NODE under the project ID OEP000957, and the experiment ID are OEX003653∼OEX003658. These data are available under from the corresponding author on reasonable request.

Meanwhile, metagenomic data from *Tara* Oceans Project used in this study are under project ID PRJEB1787, PRJEB1788, PRJEB4352, PRJEB4419. Besides, accession numbers of rhizosphere metagenomic data are provided in Supplementary Table 2. And EMP data is available on https://earthmicrobiome.org/.

## Acknowledgement

This work was supported by National Natural Science Foundation of China (81774152 to RXZ; 41676177, 91951117 to YZ; 41776173 to XX), Natural Science Foundation, the Shanghai Committee of Science and Technology 16ZR1449800 (to RXZ). The funders had no role in study design, data collection and analysis, decision to publish, or preparation of the manuscript.

## Author contribution statement

RXZ and YZ conceived and designed the project. Each author has contributed significantly to the submitted work. JX and YZ drafted the manuscript. WNC, YBX, RZ, LWT, YQW, YDZ, XX and RXZ revised the manuscript. All authors read and approved the final manuscript.

