## Supplementary for "Diversity and biogeography of Woesearchaeota: A comprehensive analysis of multi-environment data"

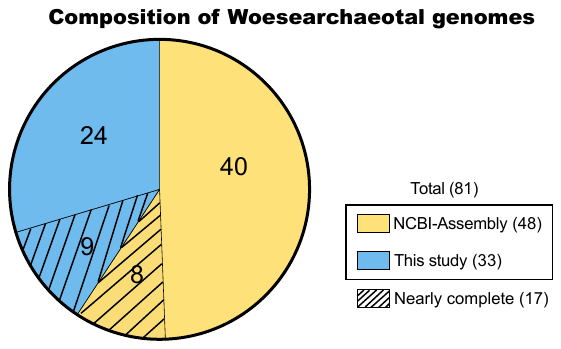

**Supplementary Fig.1 Composition of Woesearchaeotal genomes.** 81 high quality Woesearchaeotal genomes used for Taxonomy analysis are from two sources, including 48 genomes (yellow) from NCBI Assembly database, and 33 genomes reconstructed from metagenomic data in this study(blue). Meanwhile, 17 genomes are nearly complete.

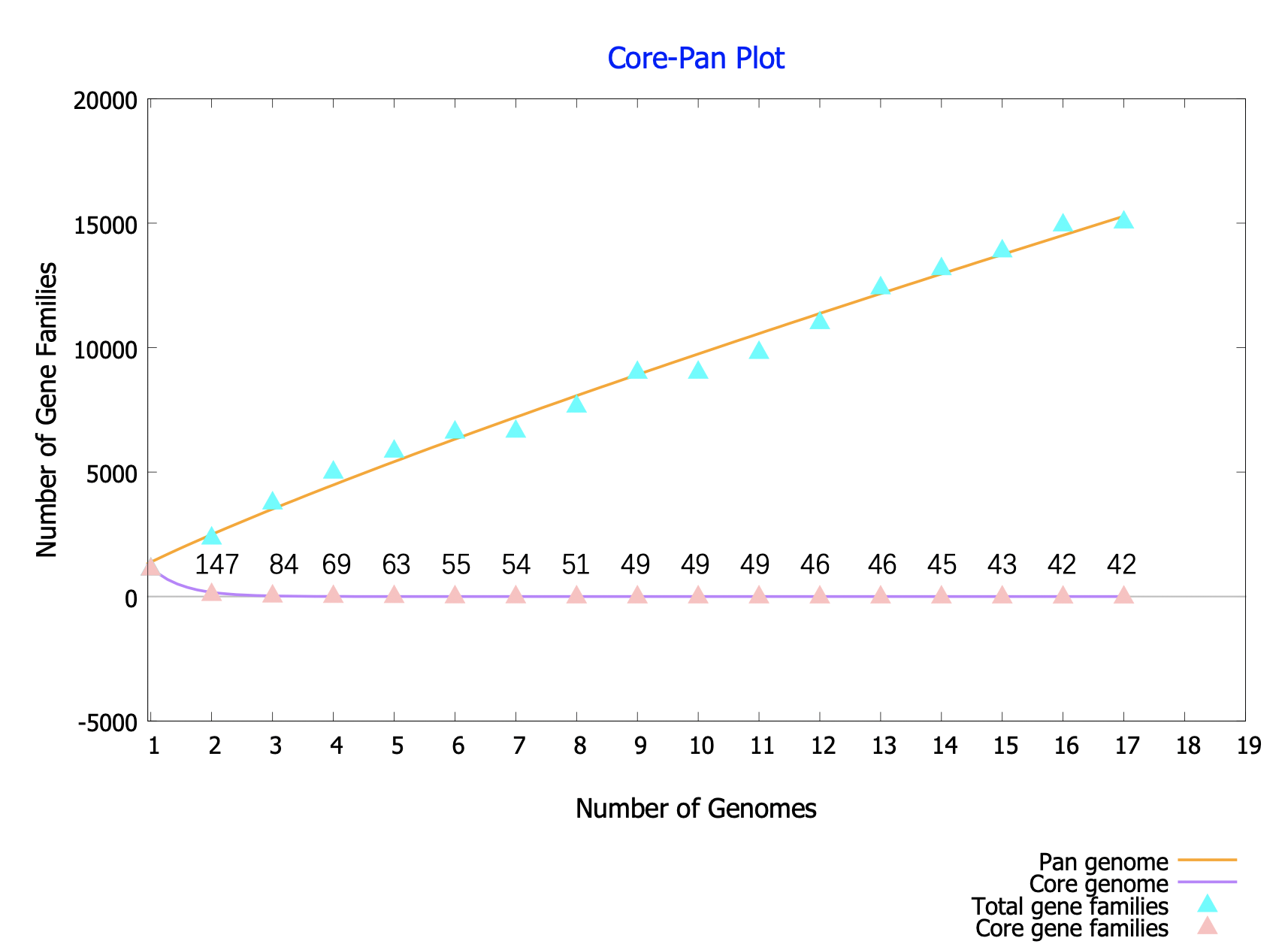

**Supplementary Fig.2 Pan and core genome** **curves of 17 Woesearchaeotal genomes.** The pan genome curve shows the trend of total number of gene families with sequential addition of more genomes, and the core-genome curve depicts the trend of shared gene families with sequential addition of more genomes. Numbers near the pink triangle represent core gene numbers.

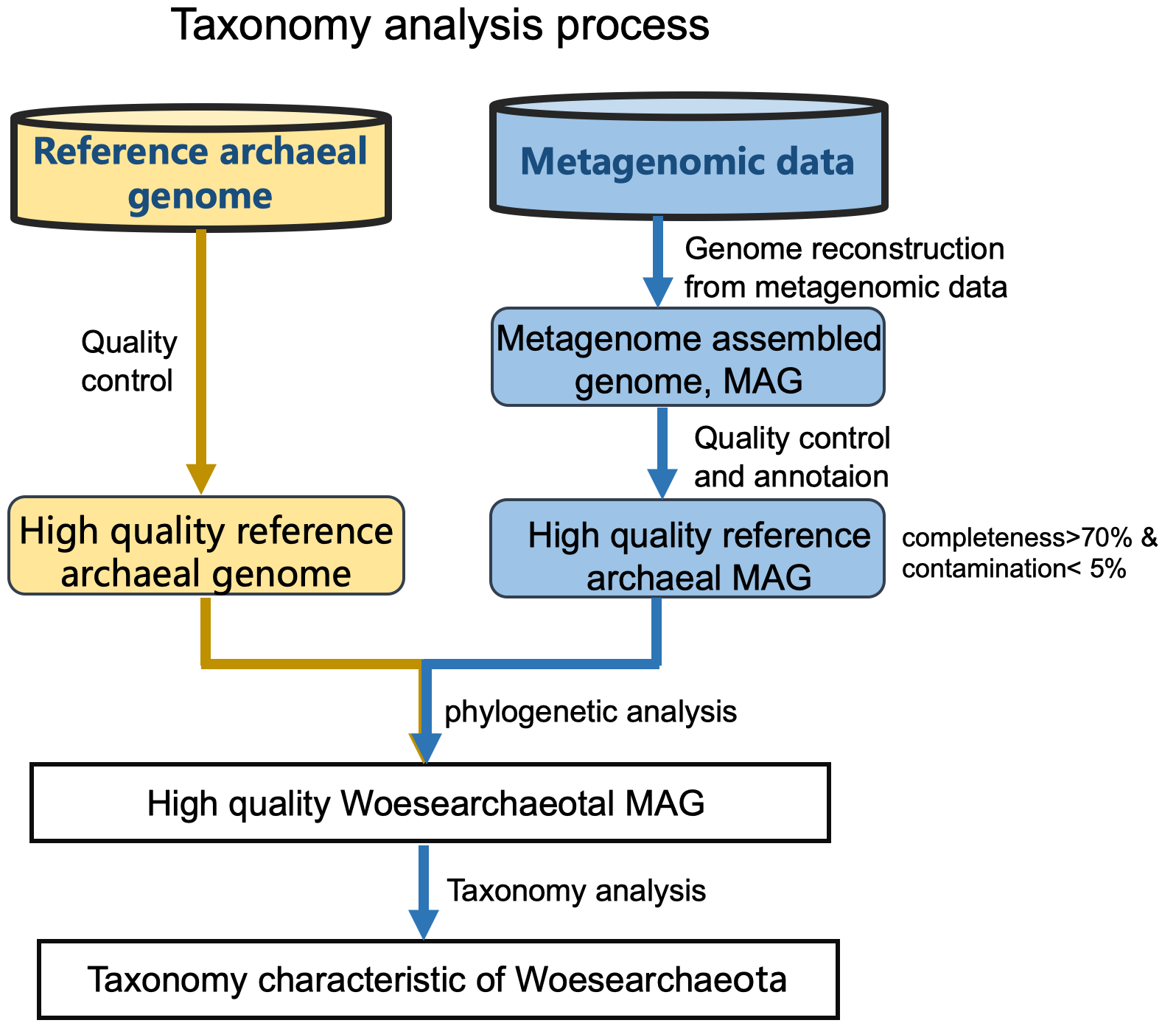

**Supplementary Fig.3** **Taxonomy analysis process.** Two types of data are used for Taxonomy analysis, including reference archaeal genomes (yellow) and metagenomic whole genome sequencing data (blue).

Supplementary Table1. 81 high quality genomes of Woesearchaeota.

| Bin id | Source | Completeness | Con  tamina  tion | Strain heterogeneity | Genome size (bp) | Contigs | N50 (contigs) | Mean contig length (bp) | GC | predicted genes |
| --- | --- | --- | --- | --- | --- | --- | --- | --- | --- | --- |
| GCA_002505845.1 | NCBI-Assembly | 87.27 | 0 | 0 | 784936 | 45 | 40014 | 17367 | 36.83 | 919 |
| GCA_007117065.1 | NCBI-Assembly | 94.23 | 0 | 0 | 907019 | 51 | 28799 | 17784 | 39.78 | 1090 |
| GCA_003561825.1 | NCBI-Assembly | 83.7 | 2.56 | 66.67 | 1106493 | 226 | 5163 | 4895 | 41.99 | 1345 |
| GCA_002686295.1 | NCBI-Assembly | 90.11 | 0 | 0 | 857738 | 51 | 17470 | 16817 | 30.16 | 940 |
| GCA_002686855.1 | NCBI-Assembly | 82.42 | 0 | 0 | 1259110 | 78 | 19079 | 16141 | 33.09 | 1319 |
| GCA_002687795.1 | NCBI-Assembly | 79.76 | 1.1 | 0 | 925456 | 39 | 28579 | 23724 | 38.27 | 1007 |
| GCA_002498125.1 | NCBI-Assembly | 95.6 | 1.1 | 0 | 1358156 | 53 | 45011 | 25595 | 56.19 | 1560 |
| GCA_002792055.1 | NCBI-Assembly | 84.62 | 1.1 | 0 | 965048 | 144 | 8147 | 6685 | 33.12 | 1158 |
| GCA_001871415.1 | NCBI-Assembly | 90.66 | 0 | 0 | 1534662 | 85 | 28595 | 18052 | 57.09 | 1516 |
| GCA_002762785.1 | NCBI-Assembly | 97.8 | 0 | 0 | 1178529 | 8 | 256322 | 147302 | 44.81 | 1300 |
| GCA_007128245.1 | NCBI-Assembly | 86.26 | 1.18 | 50 | 924936 | 111 | 9869 | 8332 | 47.23 | 1149 |
| GCA_003695265.1 | NCBI-Assembly | 96.15 | 1.1 | 0 | 1055801 | 103 | 14333 | 10250 | 40.33 | 1169 |
| GCA_002505585.1 | NCBI-Assembly | 91.76 | 0 | 0 | 766458 | 24 | 44323 | 31893 | 33.02 | 913 |
| GCA_007117735.1 | NCBI-Assembly | 86.81 | 3.3 | 0 | 690413 | 50 | 23618 | 13808 | 35.4 | 798 |
| GCA_003694805.1 | NCBI-Assembly | 90.3 | 0.25 | 0 | 1210546 | 176 | 13004 | 6878 | 51.75 | 1215 |
| GCA_003695045.1 | NCBI-Assembly | 89.38 | 1.1 | 0 | 1061362 | 159 | 7856 | 6675 | 48.17 | 1213 |
| GCA_002762985.1 | NCBI-Assembly | 96.7 | 1.1 | 0 | 1515128 | 45 | 64450 | 33660 | 44.41 | 1567 |
| GCA_003599145.1 | NCBI-Assembly | 93.13 | 0 | 0 | 1306829 | 6 | 421023 | 217804 | 37.92 | 1483 |
| GCA_007131205.1 | NCBI-Assembly | 94.51 | 0 | 0 | 930305 | 84 | 15600 | 11075 | 40.99 | 1136 |
| GCA_002506165.1 | NCBI-Assembly | 90.49 | 0 | 0 | 768923 | 39 | 35125 | 19619 | 33.03 | 913 |
| GCA_003694385.1 | NCBI-Assembly | 84.71 | 1.1 | 100 | 889737 | 130 | 8140 | 6844 | 57.03 | 1005 |
| GCA_003560545.1 | NCBI-Assembly | 81.87 | 0 | 0 | 915355 | 113 | 10535 | 8100 | 43.26 | 1111 |
| GCA_001872825.1 | NCBI-Assembly | 82.42 | 1.89 | 14.29 | 968541 | 97 | 10856 | 9982 | 47.03 | 954 |
| GCA_002762865.1 | NCBI-Assembly | 91.21 | 2.2 | 50 | 1052514 | 114 | 16881 | 9210 | 33.28 | 1227 |
| GCA_002867475.1 | NCBI-Assembly | 98.9 | 2.2 | 66.67 | 1671818 | 85 | 31020 | 19668 | 31 | 1749 |
| GCA_003694495.1 | NCBI-Assembly | 89.01 | 1.1 | 0 | 843393 | 106 | 10480 | 7956 | 36.22 | 977 |
| GCA_003599055.1 | NCBI-Assembly | 80.77 | 1.1 | 100 | 1153068 | 26 | 66701 | 44348 | 51.52 | 1323 |
| GCA_002688315.1 | NCBI-Assembly | 98.63 | 0 | 0 | 912474 | 15 | 95926 | 60830 | 36.77 | 1050 |
| GCA_002779235.1 | NCBI-Assembly | 91.76 | 1.1 | 0 | 923120 | 93 | 11036 | 9919 | 44.26 | 1006 |
| GCA_002780105.1 | NCBI-Assembly | 79.67 | 1.1 | 100 | 859843 | 111 | 9556 | 7737 | 32.95 | 1021 |
| GCA_002687275.1 | NCBI-Assembly | 85.99 | 0 | 0 | 1087504 | 55 | 27688 | 19772 | 49.67 | 1338 |
| GCA_002503705.1 | NCBI-Assembly | 91.15 | 0 | 0 | 1238951 | 111 | 16311 | 11152 | 59.63 | 1315 |
| GCA_002685855.1 | NCBI-Assembly | 84.07 | 1.1 | 0 | 953131 | 66 | 16541 | 14441 | 42.3 | 1084 |
| GCA_007116645.1 | NCBI-Assembly | 82.42 | 1.1 | 0 | 900746 | 84 | 13602 | 10723 | 44.85 | 1123 |
| GCA_902385765.1 | NCBI-Assembly | 86.12 | 0 | 0 | 1041753 | 98 | 13096 | 10630 | 42.36 | 1306 |
| GCA_002792115.1 | NCBI-Assembly | 92.86 | 0 | 0 | 1329853 | 68 | 37563 | 19549 | 33.22 | 1468 |
| GCA_005222965.1 | NCBI-Assembly | 99.18 | 1.65 | 0 | 1021708 | 4 | 1000545 | 255427 | 30.17 | 1107 |
| GCA_002762795.1 | NCBI-Assembly | 91.76 | 0 | 0 | 1351557 | 41 | 76824 | 32946 | 36.82 | 1517 |
| GCA_002762845.1 | NCBI-Assembly | 86.81 | 0 | 0 | 859335 | 91 | 14268 | 9437 | 35.94 | 932 |
| GCA_003695435.1 | NCBI-Assembly | 91.21 | 0 | 0 | 899557 | 71 | 17581 | 12669 | 41.23 | 1004 |
| GCA_002763335.1 | NCBI-Assembly | 93.13 | 0 | 0 | 1216694 | 295 | 6366 | 4110 | 46.74 | 1315 |
| GCA_002503425.1 | NCBI-Assembly | 88.83 | 0 | 0 | 1546092 | 174 | 15270 | 8835 | 57.12 | 1562 |
| GCA_002503745.1 | NCBI-Assembly | 88.74 | 0 | 0 | 830847 | 42 | 33696 | 19697 | 36.79 | 972 |
| GCA_002794135.1 | NCBI-Assembly | 82.25 | 0 | 0 | 1637751 | 280 | 7006 | 5840 | 42.58 | 1786 |
| GCA_002762705.1 | NCBI-Assembly | 87.55 | 0 | 0 | 1840814 | 183 | 12782 | 10046 | 42.51 | 1850 |
| GCA_000830315.1 | NCBI-Assembly | 85.99 | 0 | 0 | 829411 | 1 | 829408 | 829408 | 31.67 | 1015 |
| GCA_000830295.1 | NCBI-Assembly | 98.9 | 1.1 | 0 | 1157790 | 1 | 1157690 | 1157690 | 42.49 | 1309 |
| GCA_000806155.1 | NCBI-Assembly | 82.33 | 0 | 0 | 822540 | 19 | 53959 | 43289 | 29.72 | 1062 |
| 5414.119 | In-house | 84.07 | 0 | 0 | 989645 | 65 | 29992 | 15225 | 28.36 | 1090 |
| EPRCH7.246 | In-house | 86.54 | 0 | 0 | 1124680 | 88 | 18513 | 12780 | 53.83 | 1209 |
| EPRCH7.416 | In-house | 91.21 | 0.08 | 0 | 862294 | 43 | 35267 | 20053 | 37.88 | 926 |
| EPRCH7.420 | In-house | 95.6 | 1.1 | 0 | 1458275 | 41 | 52760 | 35567 | 38.83 | 1410 |
| EPRCH7.83 | In-house | 96.7 | 1.1 | 0 | 1162916 | 122 | 12704 | 9532 | 33.75 | 1195 |
| ERR594290.11 | *Tara* ocean | 91.76 | 0 | 0 | 735471 | 12 | 73490 | 61289 | 33.12 | 880 |
| ERR594290.61 | *Tara* ocean | 92.77 | 0 | 0 | 912986 | 33 | 35096 | 27666 | 34.67 | 1014 |
| ERR594290.9 | *Tara* ocean | 81.32 | 0 | 0 | 694219 | 58 | 24414 | 11969 | 32.25 | 880 |
| ERR594345.116 | *Tara* ocean | 97.53 | 0 | 0 | 1248950 | 59 | 42638 | 21168 | 35.56 | 1431 |
| ERR594345.135 | *Tara* ocean | 96.43 | 1.1 | 0 | 900222 | 29 | 39681 | 31042 | 36.74 | 1051 |
| ERR594345.138 | *Tara* ocean | 87.91 | 0 | 0 | 785881 | 41 | 37972 | 19167 | 32.17 | 963 |
| ERR594345.139 | *Tara* ocean | 89.01 | 3.3 | 0 | 749354 | 35 | 44864 | 21410 | 27.31 | 950 |
| ERR594345.145 | *Tara* ocean | 90.66 | 0 | 0 | 732537 | 18 | 71814 | 40696 | 33.13 | 883 |
| ERR594345.17 | *Tara* ocean | 84.34 | 2.38 | 75 | 1468414 | 127 | 17566 | 11562 | 38.24 | 1773 |
| ERR594345.31 | *Tara* ocean | 96.15 | 2.2 | 0 | 1378167 | 57 | 88866 | 24178 | 35 | 1593 |
| ERR594345.48 | *Tara* ocean | 96.98 | 2.2 | 33.33 | 1319769 | 90 | 29924 | 14664 | 34.9 | 1561 |
| ERR594345.81 | *Tara* ocean | 87.36 | 2.2 | 0 | 678227 | 29 | 46639 | 23387 | 26.61 | 830 |
| ERR594345.94 | *Tara* ocean | 91.67 | 0 | 0 | 917562 | 34 | 37583 | 26987 | 34.68 | 1010 |
| ERR594346.2 | *Tara* ocean | 91.76 | 2.2 | 0 | 981634 | 64 | 26960 | 15338 | 27.97 | 1083 |
| ERR594346.21 | *Tara* ocean | 88.46 | 2.2 | 0 | 776412 | 43 | 39986 | 18056 | 32.27 | 962 |
| ERR594346.48 | *Tara* ocean | 85.35 | 0 | 0 | 489031 | 21 | 42469 | 23287 | 36.14 | 612 |
| ERR594346.82 | *Tara* ocean | 91.67 | 1.1 | 0 | 777836 | 28 | 48153 | 27779 | 33.09 | 936 |
| ERR599015.28 | *Tara* ocean | 93.32 | 2.2 | 50 | 751538 | 100 | 8949 | 7515 | 33.95 | 927 |
| ERR599062.136 | *Tara* ocean | 97.53 | 1.1 | 0 | 922514 | 33 | 56027 | 27954 | 36.72 | 1075 |
| ERR599062.99 | *Tara* ocean | 89.01 | 0 | 0 | 668008 | 26 | 36304 | 25692 | 27.3 | 837 |
| ERR599076.5 | *Tara* ocean | 92.58 | 4.95 | 12.5 | 1012867 | 162 | 7313 | 6252 | 34.09 | 1220 |
| ERR599127.45 | *Tara* ocean | 87.55 | 0 | 0 | 893416 | 139 | 8279 | 6427 | 34.09 | 1080 |
| ERR599051.1 | *Tara* ocean | 78.3 | 2.2 | 0 | 636729 | 110 | 6405 | 5788 | 34.15 | 784 |
| SRR8464958.40 | NCBI-SRA | 91.91 | 0.55 | 0 | 1143662 | 86 | 19466 | 13298 | 37.87 | 1275 |
| SRR8464959.59 | NCBI-SRA | 92.86 | 2.2 | 0 | 1146943 | 55 | 31214 | 20853 | 38.67 | 1261 |
| SRR8464959.7 | NCBI-SRA | 96.89 | 0 | 0 | 1309954 | 28 | 81594 | 46784 | 31.71 | 1332 |
| SRR8464960.7 | NCBI-SRA | 95.97 | 0 | 0 | 1375015 | 41 | 62521 | 33536 | 31.68 | 1443 |
| SRR8543847.128 | NCBI-SRA | 93.04 | 1.65 | 0 | 1307620 | 109 | 18553 | 11996 | 31.7 | 1407 |

Supplementary Table 2. Metagenomic data of rhizosphere samples from NCBI-SRA.

| Run-ID | bases | size_MB | SRAStudy | BioProject | BioSample |
| --- | --- | --- | --- | --- | --- |
| SRR7975608 | 6.4465E+10 | 28549 | SRP164296 | PRJNA444376 | SAMN08779532 |
| SRR6956985 | 5.6243E+10 | 26561 | SRP137921 | PRJNA444377 | SAMN08777461 |
| SRR7975684 | 6.0375E+10 | 28702 | SRP164298 | PRJNA444378 | SAMN08779533 |
| SRR6957156 | 5.9762E+10 | 29820 | SRP137994 | PRJNA444379 | SAMN08778429 |
| SRR6957165 | 5.2202E+10 | 25257 | SRP137998 | PRJNA444380 | SAMN08776596 |
| SRR6957162 | 5.3309E+10 | 28080 | SRP137996 | PRJNA444381 | SAMN08777425 |
| SRR7975607 | 6.0946E+10 | 25849 | SRP164295 | PRJNA444382 | SAMN08779534 |
| SRR7975588 | 4.8083E+10 | 19914 | SRP164293 | PRJNA444383 | SAMN08779529 |
| SRR7975950 | 3.1374E+10 | 12521 | SRP164302 | PRJNA444384 | SAMN08779530 |
| SRR6957160 | 5.6097E+10 | 27835 | SRP137995 | PRJNA444385 | SAMN08776902 |
| SRR7975954 | 4.4733E+10 | 18339 | SRP164305 | PRJNA444386 | SAMN08779531 |
| SRR6957362 | 5.3455E+10 | 22168 | SRP138005 | PRJNA444387 | SAMN08776595 |
| SRR6435887 | 2775460030 | 926 | SRP127862 | PRJNA406144 | SAMN07632334 |
| SRR6435888 | 2881906574 | 950 | SRP127863 | PRJNA406148 | SAMN07632338 |
| SRR6435947 | 3323978704 | 1421 | SRP127867 | PRJNA406149 | SAMN07632339 |
| SRR6435948 | 4752698726 | 1800 | SRP127868 | PRJNA406150 | SAMN07632059 |
| SRR6435949 | 3197743006 | 1368 | SRP127869 | PRJNA406151 | SAMN07632060 |
| SRR6435946 | 4946114928 | 1865 | SRP127866 | PRJNA406152 | SAMN07632061 |
| SRR6435951 | 4364301056 | 1850 | SRP127871 | PRJNA406153 | SAMN07632062 |
| SRR6435956 | 2829376392 | 941 | SRP127875 | PRJNA406154 | SAMN07632063 |
| SRR6435955 | 3190537286 | 1359 | SRP127876 | PRJNA406155 | SAMN07632064 |
| SRR6435961 | 3279347634 | 1399 | SRP127880 | PRJNA406161 | SAMN07632070 |
| SRR6435958 | 2631565788 | 875 | SRP127877 | PRJNA406162 | SAMN07632071 |
| SRR6435959 | 2969194238 | 1280 | SRP127878 | PRJNA406163 | SAMN07632072 |
| SRR6435963 | 4663782074 | 1744 | SRP127882 | PRJNA406164 | SAMN07632073 |
| SRR6435960 | 4005005918 | 1698 | SRP127879 | PRJNA406165 | SAMN07632074 |
| SRR6435982 | 2491104078 | 824 | SRP127886 | PRJNA406166 | SAMN07632075 |
| SRR6435983 | 3325456088 | 1254 | SRP127887 | PRJNA406167 | SAMN07632076 |
| SRR6435984 | 4260564056 | 1602 | SRP127888 | PRJNA406168 | SAMN07632077 |
| SRR6436021 | 3159549368 | 1314 | SRP127900 | PRJNA406169 | SAMN07632078 |
| SRR6436015 | 2616418374 | 855 | SRP127894 | PRJNA406170 | SAMN07632079 |
| SRR6436020 | 3710116206 | 1573 | SRP127899 | PRJNA406171 | SAMN07632080 |
| SRR6436017 | 2687218952 | 889 | SRP127896 | PRJNA406172 | SAMN07632081 |
| SRR6436016 | 3540482202 | 1499 | SRP127895 | PRJNA406173 | SAMN07632082 |
| SRR7975952 | 3.9235E+10 | 16022 | SRP164304 | PRJNA444391 | SAMN08779515 |
| SRR6957363 | 5.9159E+10 | 28919 | SRP138006 | PRJNA444392 | SAMN08776594 |
| SRR7975957 | 5.2485E+10 | 21004 | SRP164307 | PRJNA444393 | SAMN08779527 |
| SRR6489859 | 1.4684E+10 | 6392 | SRP130755 | PRJNA406786 | SAMN07631782 |
| SRR6489884 | 1.7502E+10 | 7656 | SRP130757 | PRJNA406787 | SAMN07631783 |
| SRR6490015 | 1.8566E+10 | 7998 | SRP130767 | PRJNA406788 | SAMN07631784 |
| SRR6490096 | 2.4792E+10 | 10277 | SRP130775 | PRJNA406789 | SAMN07631785 |
| SRR7091408 | 2.4005E+10 | 10639 | SRP144277 | PRJNA441428 | SAMN08777777 |
| SRR7091409 | 2.6117E+10 | 11025 | SRP144278 | PRJNA441429 | SAMN08777778 |
| SRR7091410 | 2.7816E+10 | 11474 | SRP144279 | PRJNA441430 | SAMN08776969 |
| SRR7591305 | 2867920652 | 1256 | SRP155194 | PRJNA467686 | SAMN09201731 |
| SRR7591450 | 3234331514 | 1393 | SRP155202 | PRJNA467687 | SAMN09201123 |
| SRR7591449 | 3572630706 | 1565 | SRP155201 | PRJNA467688 | SAMN09201122 |
| SRR7591549 | 7656972628 | 3277 | SRP155208 | PRJNA467689 | SAMN09202491 |
| SRR7591534 | 6747183434 | 2943 | SRP155205 | PRJNA468122 | SAMN09202372 |
| SRR9009751 | 1.7779E+10 | 5220 | SRP195125 | PRJNA539665 | SAMN11533342 |
| SRR9009750 | 2.6588E+10 | 7766 | SRP195124 | PRJNA539666 | SAMN11532184 |
| SRR8543847 | 4.1533E+10 | 11447 | SRP184460 | PRJNA518430 | SAMN10863918 |
| SRR8543849 | 7.3857E+10 | 21673 | SRP184462 | PRJNA518431 | SAMN10864298 |
| SRR8464958 | 1.5747E+10 | 4609 | SRP180571 | PRJNA502707 | SAMN10350727 |
| SRR8464957 | 1.286E+10 | 3771 | SRP180570 | PRJNA502708 | SAMN10350572 |
| SRR8543846 | 1.9943E+10 | 5493 | SRP184459 | PRJNA518432 | SAMN10864359 |
| SRR8464959 | 1.2265E+10 | 3617 | SRP180572 | PRJNA502709 | SAMN10350739 |
| SRR8464960 | 1.3844E+10 | 4098 | SRP180573 | PRJNA502710 | SAMN10350571 |
| SRR8543848 | 1.4753E+10 | 4328 | SRP184461 | PRJNA518433 | SAMN10863917 |
| SRR8543844 | 1.9387E+10 | 5321 | SRP184457 | PRJNA518434 | SAMN10863916 |
| SRR8544061 | 5.2895E+10 | 15163 | SRP184474 | PRJNA518440 | SAMN10863913 |
| SRR8544180 | 5.3162E+10 | 15547 | SRP184479 | PRJNA518441 | SAMN10864233 |
| SRR8554882 | 6.6771E+10 | 18979 | SRP185317 | PRJNA518255 | SAMN10864097 |
| SRR9021703 | 6.3323E+10 | 17981 | SRP195735 | PRJNA539583 | SAMN11533248 |
| SRR9021705 | 5.3481E+10 | 15203 | SRP195737 | PRJNA539584 | SAMN11532178 |
| SRR8554884 | 8.2391E+10 | 23496 | SRP185319 | PRJNA518256 | SAMN10864380 |
| SRR9021691 | 6.524E+10 | 18573 | SRP195734 | PRJNA539586 | SAMN11532355 |
| SRR9021706 | 6.7821E+10 | 19421 | SRP195738 | PRJNA539587 | SAMN11532334 |
| SRR9021707 | 7.6371E+10 | 21780 | SRP195739 | PRJNA539588 | SAMN11532792 |
| SRR9021711 | 6.7279E+10 | 19270 | SRP195741 | PRJNA539589 | SAMN11532865 |
| SRR9021708 | 6.9524E+10 | 19946 | SRP195740 | PRJNA539590 | SAMN11532177 |
| SRR9045292 | 4.7529E+10 | 14113 | SRP197978 | PRJNA539071 | SAMN11532919 |
| SRR6214544 | 5.4501E+10 | 32877 | SRP121388 | PRJNA366149 | SAMN06268382 |
| SRR5508097 | 6.5545E+10 | 27871 | SRP106465 | PRJNA375567 | SAMN06344173 |
| SRR5509277 | 6.0379E+10 | 25083 | SRP106478 | PRJNA375568 | SAMN06344174 |
| SRR5511004 | 6.1944E+10 | 26163 | SRP106483 | PRJNA375569 | SAMN06344175 |
| SRR5511291 | 6.7466E+10 | 30912 | SRP106493 | PRJNA375570 | SAMN06344176 |
| SRR5511879 | 6.5953E+10 | 29868 | SRP106496 | PRJNA375571 | SAMN06344177 |
| SRR5578835 | 6.7852E+10 | 31227 | SRP107494 | PRJNA375572 | SAMN06344178 |
| SRR5578837 | 6.6432E+10 | 30335 | SRP107495 | PRJNA375573 | SAMN06344179 |
| SRR5517431 | 6.5185E+10 | 29625 | SRP106648 | PRJNA375574 | SAMN06344180 |
| SRR5517144 | 5.5824E+10 | 22906 | SRP106645 | PRJNA375575 | SAMN06344181 |
| SRR5574089 | 4.1855E+10 | 17216 | SRP107245 | PRJNA375576 | SAMN06344182 |
| SRR5576206 | 6.1682E+10 | 27680 | SRP107359 | PRJNA375577 | SAMN06344183 |
| SRR5574261 | 5.9973E+10 | 26775 | SRP107252 | PRJNA375578 | SAMN06344184 |
| SRR5574262 | 5.2032E+10 | 23795 | SRP107253 | PRJNA375579 | SAMN06344185 |
| SRR5574263 | 4.0724E+10 | 18424 | SRP107254 | PRJNA375580 | SAMN06344186 |
| SRR5579275 | 5.3298E+10 | 24587 | SRP107557 | PRJNA365487 | SAMN06266402 |
| SRR5581992 | 4.1359E+10 | 18731 | SRP107756 | PRJNA375656 | SAMN06343958 |
| SRR5582001 | 4.4254E+10 | 20001 | SRP107758 | PRJNA375657 | SAMN06343959 |
| SRR5582436 | 4.7939E+10 | 20781 | SRP107771 | PRJNA375658 | SAMN06343960 |
| SRR5582372 | 2.8463E+10 | 11792 | SRP107769 | PRJNA375659 | SAMN06343961 |
| SRR5582390 | 4.7006E+10 | 19051 | SRP107770 | PRJNA375660 | SAMN06343962 |
| SRR5582993 | 6.5442E+10 | 27832 | SRP107779 | PRJNA375661 | SAMN06343963 |
| SRR5583252 | 3.3193E+10 | 14088 | SRP107786 | PRJNA375662 | SAMN06343964 |
| SRR5583253 | 2296522458 | 1051 | SRP107786 | PRJNA375662 | SAMN06343964 |
| SRR5583254 | 6.5268E+10 | 27720 | SRP107786 | PRJNA375662 | SAMN06343964 |
| SRR5584097 | 5.573E+10 | 26354 | SRP107797 | PRJNA375663 | SAMN06343965 |
| SRR5584873 | 6.1565E+10 | 25316 | SRP107804 | PRJNA375664 | SAMN06343966 |
| SRR5585385 | 6.8544E+10 | 30062 | SRP107807 | PRJNA375665 | SAMN06343967 |
| SRR5585387 | 4.0372E+10 | 17254 | SRP107808 | PRJNA375666 | SAMN06343968 |
| SRR5585388 | 5.8476E+10 | 25187 | SRP107809 | PRJNA375667 | SAMN06343969 |
